# Light-sheet scattering microscopy to visualize long-term interactions between cells and extracellular matrix

**DOI:** 10.1101/2021.10.22.465506

**Authors:** Xiangda Zhou, Renping Zhao, Archana K. Yanamandra, Markus Hoth, Bin Qu

## Abstract

Visualizing interactions between cells and the extracellular matrix (ECM) mesh is important to understand cell behavior and regulatory mechanisms by the extracellular environment. However, long term visualization of three-dimensional (3D) matrix structures remains challenging mainly due to photobleaching or blind spots perpendicular to the imaging plane. Here, we combine label-free light-sheet scattering microcopy (LSSM) and fluorescence microscopy to solve these problems. We verified that LSSM can reliably visualize structures of collagen matrices from different origin including bovine, human and rat tail. The quality and intensity of collagen structure images acquired by LSSM did not decline with time. LSSM offers abundant wavelength choice to visualize matrix structures, maximizing combination possibilities with fluorescently-labelled cells, allowing visualizing of long-term ECM-cell interactions in 3D. Interestingly, we observed ultrathin thread-like structures between cells and matrix using LSSM, which were not observed by normal fluorescence microscopy. Transient local alignment of matrix by cell-applied forces can be observed. In summary, LSSM provides a powerful and robust approach to investigate the complex interplay between cells and ECM.

## Introduction

Under physiological conditions, immune cells and tumor cells encounter complex and dynamic three-dimensional (3D) environments. To maintain 3D environments, extracellular matrix (ECM) composed of fibrous mesh networks serves as one main structural component. Concerning immune surveillance, interaction of ECM with immune cells plays an essential role in tuning physiological responses [1]. Evidence from *in vitro* and *in vivo* shows that there are channels/tunnels in ECM, which can be either created or expanded by immune cells, facilitating migration of subsequent immune cells [2, 3]. These channels could also contribute to metastasis of tumor cells [4]. In case the channels in ECM are too narrow, the width could serve as a speed limiting factor to restrain the migration of immune cells [5]. Thus, migration of cells in ECM per se results in structural changes in ECM [2, 3]. Such structural changes may result in different outcomes of immune surveillance. To investigate this, long-term visualization of ECM-cell interaction with defined and controllable features is needed. However, long-term visualization ECM structures remains technically very challenging. One of the most applied approaches is to label the matrix protein of interest fluorescently [6]. Unfortunately, due to photobleaching, this method does not offer a satisfactory solution for long-term visualization. There are numerous other approaches well established to visualize fibrillar collagen, such as electron microscopy and second hormonic generation microscopy [7, 8]. Although these methods enable visualization of matrix structure with a high resolution, they cannot be applied for live cell imaging. A recently emerged method called confocal reflection microscopy utilizes the light reflected by the fibrous structure of the matrix [9], offering a possibility to visualize collagen structure without fluorescence, which can also be combined with long-term live cell imaging. However, with confocal reflection microscopy, the subset of fibrous structures perpendicular to the imaging plane cannot be detected due to a blind spot, resulting in incomplete information in 3D structure of the reconstructed networks [10]. Light can be scattered during its propagation in the sample in an inelastic or elastic manner. Inelastic scattering occurs when the light is scattered off at molecular bonds, resulting in change in energy and the corresponding frequency [11]. The spectrum of inelastic scattering is utilized to characterize chemical composition of samples in 3D, as for example performed with Raman scattering [12]. In contrast, when the light is scattered by spatial structures, there is no energy loss in the scattered light. This process is termed as elastic scattering. Several recent studies have used elastically scattered light with light-sheet microscopy to visualize plant roots in transparent soil or gel [13], freshly-excised tissues *ex vivo* [14] or blood cells *in vivo* [15] without fluorescent labeling.

In this paper, we verified the label-free light-sheet scattering microscopy (LSSM) as a robust approach to visualize ECM structures without blind spot and photobleaching. Non-fluorescently labeled cells can be also visualized by LSSM. LSSM is particularly advantageous to visualize long-term cell-ECM interactions, and to analyze cell behavior adapted to dynamic ECM meshworks. Interestingly, applying LSSM we observed thin thread-like structure between cells and matrix, which could not be observed by normal fluorescence microscopy. Furthermore, LSSM can be used to characterize forces exerted on the matrix by the embedded cells.

## Materials and methods

### Reagents

The following reagents were used in this work: Atto Fluor 647 NHS-Ester (Succinimidylester), Atto Fluor 488 NHS-Ester (Succinimidylester), CellTrace CFSE Cell Proliferation Kit (ThermoFisher Scientific), and CellTrace Calcein Red-Orange AM (ThermoFisher Scientific). RatCol Rat Tail Collagen (4 mg/ml), FibriCol Type I Collagen Solution (bovine, 10 mg/ml) and VitroCol Type I Collagen Solution (Human, 3 mg/ml) were obtained from Advanced BioMatrix.

### Cell preparation and cell culture

Human primary CD4^+^ T cells were negatively isolated using CD4 T Cell Isolation Kit Human (Miltenyi) from peripheral blood mononuclear cells (PBMCs) of healthy donors [16]. CD4^+^ T cells were stimulated with Dynabeads Human T-Activator CD3/CD28 (ThermoFisher Scientific) and cultured in AIM V Medium (ThermoFisher Scientific) with 10% FCS (ThermoFisher Scientific) at a density of 2×10^6^ cells/ml. Human pancreatic beta cells 1.4E7 were purchased from Merck and cultured in RPMI-1640 medium (ThermoFisher Scientific) supplemented with 2 mM glutamine, 1% Penicillin-Streptomycin plus 10% FCS. SK-MEL-5 cells were cultured in MEM medium (ThermoFisher Scientific) containing 10% FCS and 1% penicillin-streptomycin. All cells were cultured at 37 °C with 5% CO_2_.

### Preparation of collagen matrices

Collagen solutions were prepared as previously described [17]. Briefly, chilled 10× PBS was added to collagen stock solution prior to neutralization with 0.1 M NaOH. The neutralized collagen solution was further diluted to the desired concentrations with 1× PBS for structure visualization or with AIMV medium for live cell imaging. The collagen was then sucked into a glass capillary and kept at 37°C with 5% CO_2_ for 40 min for polymerization.

### Fluorescent labeling of collagen fibres

After polymerization, the collagen rod was pushed out of the capillary carefully and immersed in the freshly made Atto Fluor 647 NHS-Ester or Atto Fluor 488 NHS-Ester solution (50 μg/ml) for 10 min at room temperature, followed by three times washing with 1× PBS for 5min. For live cell imaging, additional steps were required as previously reported [6]. Briefly, the Atto Fluor 488 NHS-ester-stained collagen was dissolved in acetic acid (20 mM) at 4°C. Then the acid-dissolved fluorescently labeled collagen was mixed with non-labeled collagen in a ratio between 1:25-1:50. The cell pellet was resuspended in this mixed collagen solution and polymerization was conducted as describe above.

### Light-sheet microscopy

A Zeiss Z1 Light-sheet microscope was used for all experiments. The imaging chamber was assembled according to the manufacturer’s instructions. For LSSM, the combination of 445 nm laser (power 0.2 %), LBF 405/488/561/640, SBS LP 490-BP 420-470 with an exposure time of 30 ms was used if not mentioned otherwise. The samples were illuminated from one side if not mentioned otherwise. For fluorescence imaging modality, the following settings were used: 561 nm laser (power 0.5 %), LBF 405/488/561/640, SBS LP 560-LP 585 with an exposure time of 100 ms for calcein red-orange; 488 nm laser (power 1.0 %), LBF 405/488/561/640, SBS LP 490-BP 505-545, with an exposure time of 30 ms was used for Atto 488; 638 nm laser, LBF 405/488/561/640, SBS LP 560-LP 660 was used for Atto 647; 488 nm laser (power 0.5 %), LBF 405/488/561/640, SBS LP 510-BP 525-565 with an exposure time of 80 ms for CFSE. Time lapse with Z-stacks (step-size: 0.4-1.5 μm as indicated in the figure legends) was conducted with a 20× objective at 37°C. The intervals and total length of visualization are stated in the corresponding figure legend.

For live cell imaging, the cells were loaded with calcein red-orange (50 μM) at 37°C for 30 min or with CFSE (50 μM) at room temperature for 15 min. CFSE-stained cells were recovered in the incubator for 24 hours prior to imaging. The cells were centrifuged, and the pellet was resuspended in the collagen solution followed by 40 min-polymerization and 1 hour recovery in the incubator. 3D reconstruction, maximum intensity projection (MIP) and channel merge were generated with Zen (Zeiss). Tracking of collagen matrix displacement was manually carried out with the ImageJ plugin MTrackJ as reported elsewhere [18]. Colocalization was analyzed using ImageJ plugin JACoP (Just Another Colocalization Plugin) [19].

## Results

### LSSM is a label-free robust method to visualize collage structure

In order to visualize long-term cell-ECM interactions in larger 3D structures, we looked for a label-free method based on light-sheet microscopy. We therefore investigated whether elastically scattered light can be used to visualize ECM structures. To this end, we prepared type I bovine collagen samples in glass capillaries and visualized the collagen sample with various lasers (405 nm, 445 nm, 488 nm, 515 nm, 561 nm, 638 nm) in combination with different block filters, beam splitters and detection filters as summarized in Supplementary Fig. 1A. The acquired images show that among 216 possible combinations, the collagen structure were seen in 35 combinations (Supplementary Fig. 1B), which share a common feature: the combination of the block filter, the beam splitter and the detection filter does not exclude the wavelength of the excitation laser. In comparison, many of these working combinations do exclude theoretical emission wavelength of the auto-fluorescence, which could be excited by the corresponding laser. Out findings indicate that the structure of matrices are visualized by the elastically scattered light. This technique is hereinafter referred to as light-sheet scattering microscopy (LSSM). Of note, during acquisition, a very low laser power (0.2%) and short exposure time (150 ms) were applied, suggesting that photocytotoxicity induced by LSSM is very low if not negligible, and that a fast scan speed can be achieved.

To verify whether the matrix structure acquired by LSSM reflects the original structure of the matrix, we fluorescently labeled human collagen and scanned the same samples using LSSM and fluorescence microscopy concurrently. The results show that the fibrous structures and reconstructed 3D structures visualized by LSSM and fluorescence microscopy are very similar (Fig. 1A, Movie 1-3). Manders’ coefficients are 0.756±0.051 for M1 (scattering over fluorescence) and 0.725±0.068 for M2 (fluorescence over scattering). To examine the robustness of LSSM, we further analyzed the collagen from different origin such as bovine (Fig. 1B) and rat tail (Fig. 1C). Manders’ coefficients are 0.689±0.038 (M1) and 0.643±0.022 (M2) for bovine samples, 0.577±0.129 (M1) and 0.696±0.064 (M2) for rat tail samples. The results show that LSSM reliably obtained the key structural features of collagen matrices from different sources, with different concentrations and pore sizes.

**Figure 1.**
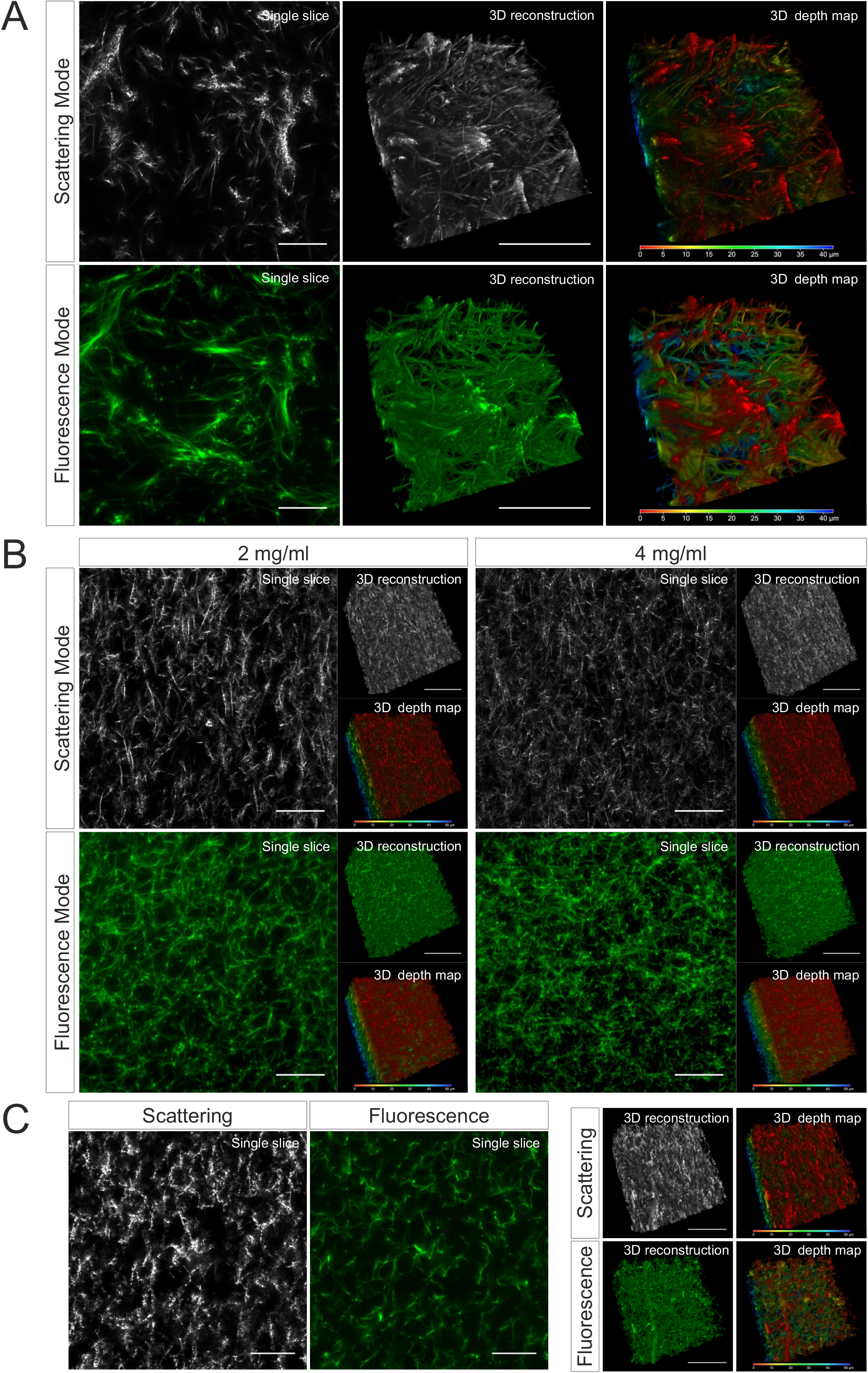
Collagen structures visualized by LSSM. Collagen from human (**A**, 2mg/ml), bovine (**B**, 2 and 4 mg/ml) or rat tail (**C**, mg/ml) was used. Collagen was fluorescently labeled with Atto 488. Z-stacks with step-size of 0.4 μm for 95 slices (**A**) or for 119 slices (**B** and **C**) were acquired. Scale bars are 40 μm. Representative images from at least three independent experiments are shown.

### No blind spot and no photobleaching in LSSM

In confocal reflection microscopy, a blind spot has been reported, where around half of the fibers, which are vertically oriented above 50 degrees from the focal plane, missing in image data acquired by confocal reflection imaging modality [20]. We therefore explored whether LSSM has the same limitation. For LSSM, we chose dual side fusion mode, in which the samples were illuminated from both the left and the right side sequentially and the images from both sides were merged to generate the final image (Fig. 2A). We noticed that although not many, a few fibers did appear in images from only one side (Fig. 2B, Scattering Mode, insets 1-2), indicating that in LSSM modality illumination from only one side could leave a few fibers undetected, presumably due to the orientation of the fibers. Interestingly, a few fibrous structures were detected only in LSSM (Fig. 2B, inset 1, comparing Scattering Mode and Fluorescence Mode). This implies that for thin fibrous structures, LSSM could be advantageous compared to fluorescence microscopy.

**Figure 2.**
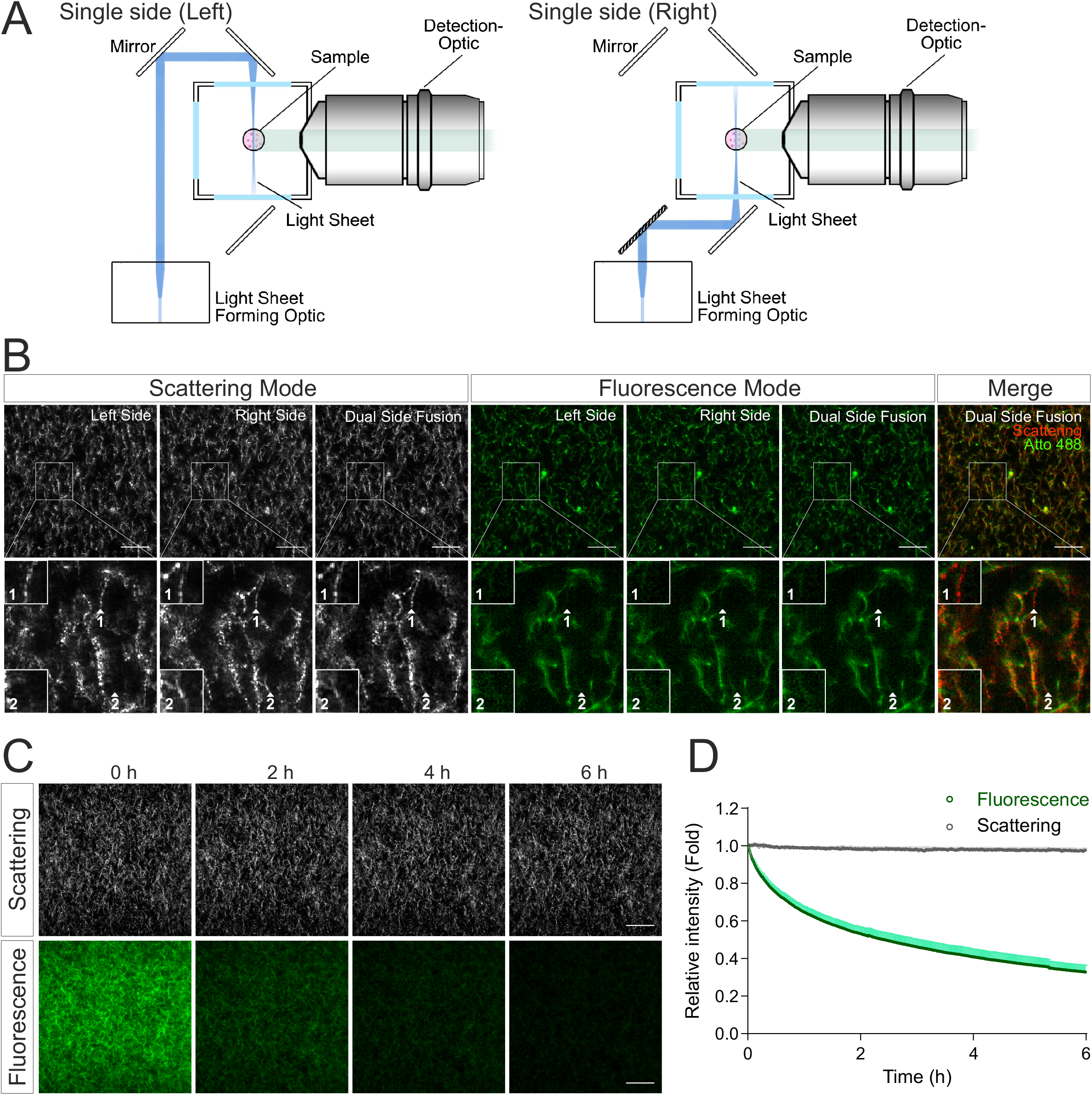
LSSM avoids photobleaching and blind spot problems. (**A**) Sketch for single side illumination. (**B**) Comparison of images acquired by LSSM and fluorescence microscopy. Rat tail collagen (2 mg/ml) was fluorescently labeled with Atto 488. Images from single side illumination and the merged images of both sides (dual side fusion) are shown. Z-stacks with step-size of 1 μm for 51 slices were obtained with LSSM and fluorescence mode concurrently at 37°C. (**C**, **D**) Signal intensity obtained by LSSM is not reduced with time. Bovine collagen (2 mg/ml) was labeled with Atto 647. Z-stacks with step-size of 0.621 μm for 50 slices were obtained with LSSM and fluorescence mode subsequentially at 37°C every 30 sec for 6 hours. Scale bars are 40 μm. Representative images from at least three independent experiments are shown.

Photobleaching is a major bottle neck for long-term visualization, at least for fluorescence microscopy. We next analyzed photobleaching in LSSM. Images of fluorescently labeled collagen matrices were acquired using LSSM and fluorescence imaging modality every 30 sec for 6 hours. Time lapse images and the analysis show a substantial photobleaching in fluorescence microscopy, while no photobleaching was detected in LSSM (Fig. 2C, D). This technical advantage of LSSM enables reliable long-term visualization of dynamic change of matrix structures as well as matrix-cell interaction.

### Application of LSSM to investigate cell-ECM interaction

Characterization of ECM-cell interaction is essential to understand the behavior of cells in 3D. We examined whether LSSM could be applied to meet this purpose. We first loaded SK-MEL-5 cells, a melanoma cell line, with calcein red-orange and embedded the cells in the Atto 488-labeled collagen matrix. Z-stacks of the samples were taken at 37°C every 30 sec for about half an hour. Interestingly, in LSSM, the SK-MEL-5 cells were also detected, and the could be morphologically distinguished from the mesh network of collagen matrix (Fig. 3A). Unexpectedly, highly dynamic filamentous structures linking the cells and the mesh network were detected in LSSM, which could not be detected by fluorescence modality (Fig. 3A, Movie 4, highlighted by the arrowheads), implying that those structures do not originate from the collagen matrix but rather from the SK-MEL-5 cells. Time lapse images from LSSM show that these dynamic filamentous structures were formed between SK-MEL-5 cells and the matrix when the cells moved away from the matrix (Fig. 3A, 7.5 min and 8 min, Movie 4), and also formed between SK-MEL-5 cells when the two cells migrated apart (Fig. 3A, 20 min and 20.5 min, Movie 4).

**Figure 3.**
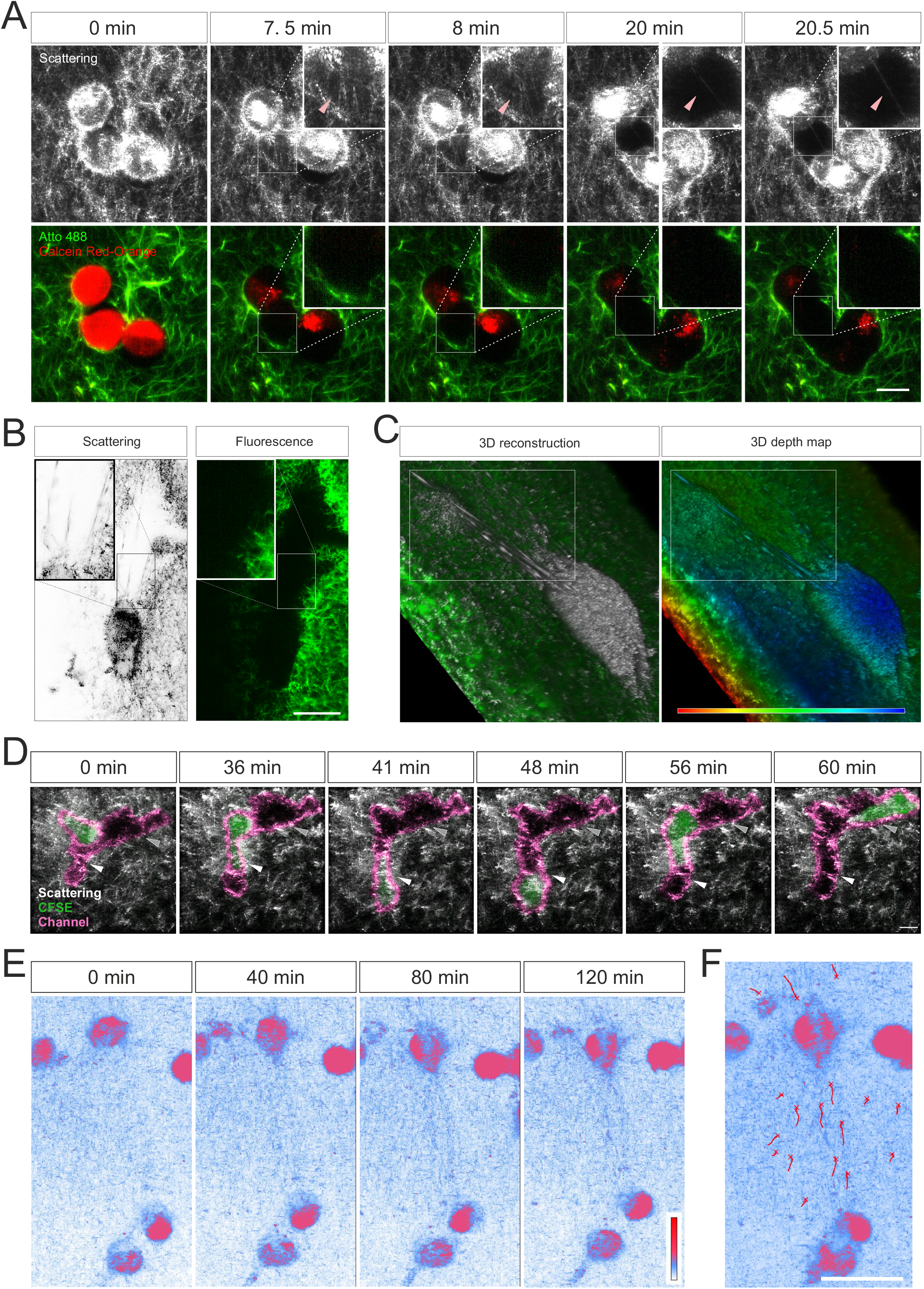
Live cell imaging with LSSM. (**A**-**C**) Ultrathin filamentous structures observed with LSSM. Rat tail collagen (2 mg/ml) was fluorescently labeled with Atto 488. Calcein red-orange loaded SK-MEL-5 cells (**A**) or non-labeled 1.4E7 human pancreatic beta cells (**B**, **C**) were embedded in the matrix. Z-stacks with step-size of 0.82 μm for 52 slices (**A**) or 0.418 μm for 82 slices (**B**, **C**) were obtained using LSSM and fluorescence modality concurrently at 37°C every 30 sec for 35.5 min (**A**) or every 20 sec for 10 min (**B**, **C**). (**D**) T cells enter channels in ECM. Primary human CD4^+^ T cells were loaded with CFSE and were embedded in the collagen matrices (Bovine collagen, 2 mg/ml). Z-stacks with step-size of 1.5 μm for 70 slices were obtained using LSSM and fluorescence modality concurrently at 37°C every 1 min for 1 hour. (**E**, **F**) Cell-induced displacement of collagen fibers. Non-labeled SK-MEL-5 cells were embedded in rat tail collagen (2 mg/ml). Z-stacks with step-size of 1 μm for 295 slices were obtained using LSSM at 37°C every 1 min for 2 hours. Scale bars are 20 μm. Representative events from three independent experiments are shown.

To identify whether these filamentous structures are unique to SK-MEL-5 cells, we used 1.4E7 cells, a human pancreatic beta cell line. Again, highly dynamic ultrathin filamentous structures between the cells and the mesh network were observed in LSSM, which could not be detected by fluorescence modality (Fig. 3B inset, Movie 5 highlighted in the upper frame). In addition, distortion of the matrix caused by cell contraction was also observed in both imaging modalities (Movie 5, highlighted in the lower frame). 3D reconstructions and the depth map (pseudocolors are assigned to different depth) further reveal that those filamentous structures were parallelly orientated in 3D and could be traced back to the cell body (Fig. 3C). These findings indicate that these LSSM-identified filamentous structures could play an essential role in regulating ECM-cell interaction and cell-cell interaction, especially in 3D environments.

We examined whether LSSM is suitable to visualize fast migrating immune cells and their interaction with ECM. We loaded primary human CD4^+^ T cells with CFSE, and visualized T cell migration in label-free 3D collagen matrices using LSSM in combination with fluorescence microscopy. Channels in collagen matrices were detected in LRSM as reported previously using fluorescence microscopy [2]. We observed that after entry into the channel, in this example the CD4^+^ T cell first chose the segment with a narrower width (Fig. 3D, highlighted by the white arrowhead, Movie 6), expanded the channel and then reversed the direction and advanced to the other segmentation (Fig. 3D, highlighted by the grey arrow head, Movie 6). That result shows that LSSM is a powerful tool to characterize matrix structure-regulated migration behavior.

Mechanical forces are an essential factor to modulate cell function and the interplay between ECM and cells. When cells exert mechanical forces on the matrix, deformation of matrix is induced and can be used to characterize the mechanical forces exerted by the cells [21, 22]. Using LSSM, this deformation could be observed as shown in Movie 5. We further examined whether this deformation could be characterized with LSSM in long-term measurements. To this end, we embedded not-fluorescently labeled SK-MEL-5 cells in collagen matrices and acquired images every 1 min for 2 hours. Time lapse images show that with LSSM, SK-MEL-5 cells can be distinguished from the ECM based on their morphology. Interestingly, collagen fibers between two SK-MEL-5 cells were reorientated and aligned towards the cells at a macroscopic length scale (Fig. 3E, F, Movie 7). These results indicate that LSSM can be applied to characterize cell-applied mechanical forces.

## Discussion

In this work, we report that ECM meshwork can be reliably visualized by LSSM. One major advantage of LSSM is that the signal intensity does not decline over time, which makes LSSM a powerful tool for long-term measurements. Another advantage of LSSM is that in contrast to reflection confocal microscopy, there is no blind spot, allowing label-free reconstruction of 3D ECM structures without discontinuous artefacts. In addition, LSSM can detect thin fibrous structures generated by cells. For ECM-cell interaction, this robust method can be combined with fluorescence modality to reveal more details of cell-ECM interactions.

LSSM utilizes the light scattered sidewards by the collagen fibers, which is detected by the objective perpendicular to the imaging plane. Here, the imaging plane is illuminated by a thin sheet of light, the incidence angle of which is parallel to the imaging plane. In comparison, images taken by confocal reflection microscopy are formed from the light reflected by the collagen fibers. Therefore, the fibers vertically orientated to the imaging plane are not detected, leading to an artefact that 3D reconstructed matrix structures seem to be aligned to the imaging plane. For LSSM, however, the possibility for a total reflection by a fiber leading to a blind spot is theoretically extremely low. As shown in our results, the fibrous structures oriented to all directions were observed, and no discontinuation of vertically oriented fibers in 3D reconstructed networks were detected. In addition, since LSSM uses the scattered light, the signal intensity is not reduced with time. Thus, LSSM is shown to be a reliable label-free method to visualize mesh networks without photobleaching, which is especially advantageous for long-term measurements.

Using LSSM, we unexpectedly observed ultrathin filamentous structures between diverging cells or between matrices and the moving cells. Since these structures were not detected in fluorescence microscopy and the collagen was fluorescently labelled, the ultrathin filamentous structures are likely to originate from the cells. This postulation is further supported by our observation that these structures appear to be generated between diverging cells. These ultrathin structures could be derived from plasma membrane with or without architectural support of cytoskeleton. Several types of thin cell protrusions are reported such as invadopodia and microtentacles. Invadopodia are thin, short, actin-rich structures, which are formed to protrude into the ECM [23]. Microtentacles are needle-like structures formed by microtubules and intermediate filaments, which are found in many tumor cells, especially in circulating tumor cells, suggestively contributing to cancer metastasis [24]. Whether the origin, generation mechanism and functions of the thin structures detected by LSSM share common features with invadopodia or microtentacles require further investigation. Mechanical interplay between cells and matrix plays a pivotal role for cell migration, differentiation, self-organization, and activation of cells [21, 22, 25–27]. Cell-cell distant mechanical communications via matrix is essential for cell self-assembly [22]. Traction forces generated by cells lead to matrix deformation, enabling force transmission to distant cells. In a 3D scenario, traction forces generated by a single cell or spheroid can be well determined by traction force microscopy, which measures the displacement of microbeads evenly distributed in the matrix [28, 29]. We observed that collagen fibers were aligned between two SK-MEL-5 cells. Emerging evidence shows that aligned matrix potentiates cell migration to a fast mode [26]. Tracking migrating immune cells in 3D matrices using LSSM can address the question that whether this cell-induced transient local alignment of matrix could be used by immune cells as directional cues to quickly find their target cells.

In summary, we present LSSM as a reliable and robust tool for long-term visualization of ECM-cell interactions. Using this assay, important scientific questions can be addressed. These include measurements of cell-cell mechanical interactions, of long-range traction force transmission, and of dynamic ECM-immune cell interactions, which may change migration patterns or even search strategies.

## Acknowledgements

We thank the Institute for Clinical Hemostaseology and Transfusion Medicine for providing donor blood; Carmen Hässig, Cora Hoxha, Sandra Janku, and Gertrud Schäfer for excellent technical help. This project was funded by the Deutsche Forschungsgemeinschaft (SFB 1027 Project A2 to BQ, and GZ: INST 256/419-1 FUGG to MH), and by Bundesministerium für Bildung und Forschung (031L0133 to MH). The authors declare no competing financial interests.

## Ethical considerations

Research carried out for this study with healthy donor material (leukocyte reduction system chambers from human blood donors) is authorized by the local ethic committee (84/15; Prof. Dr. Rettig-Stürmer).

