## Supplementary Information for "Light-sheet scattering microscopy to visualize long-term interactions between cells and extracellular matrix"

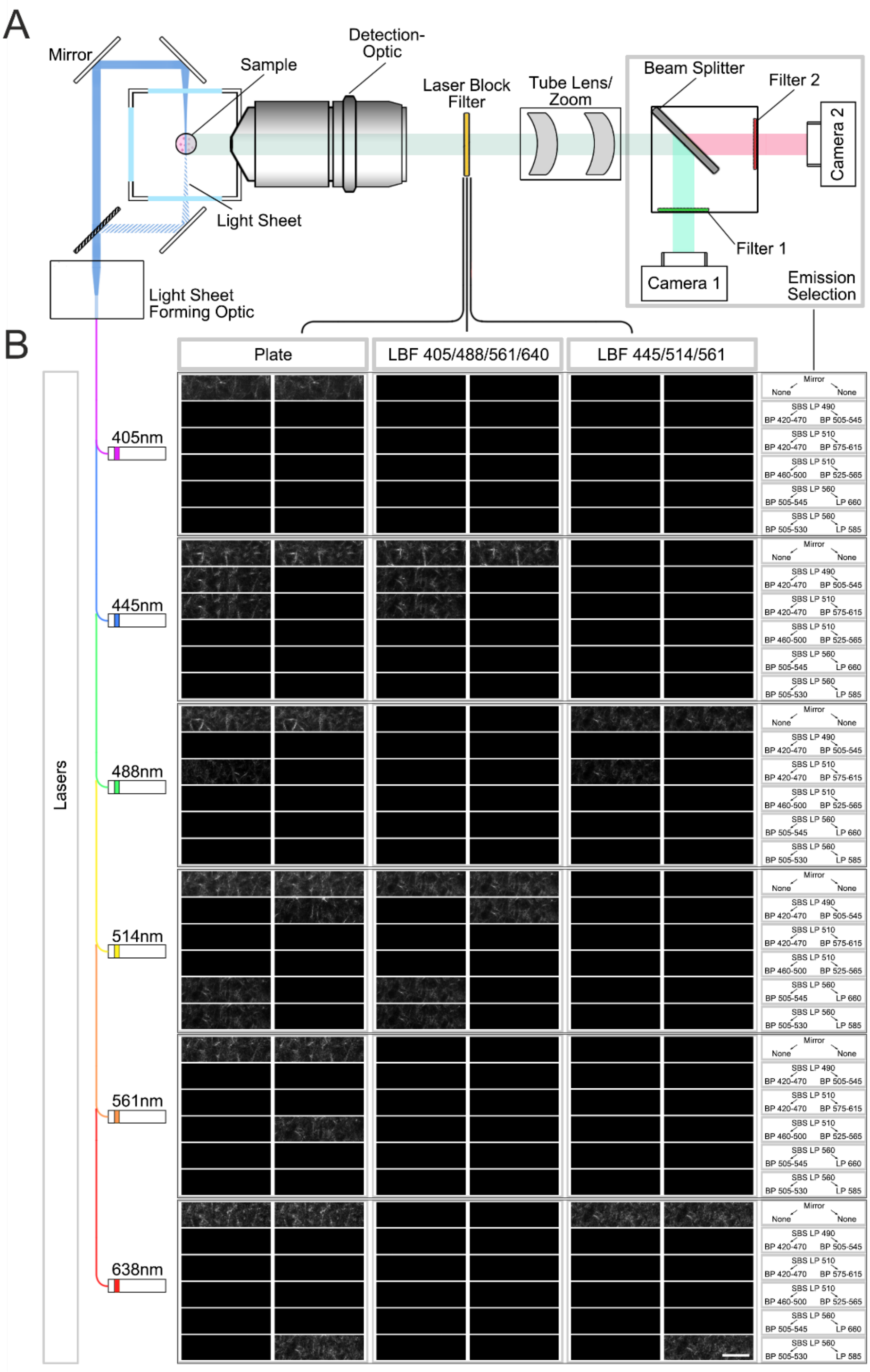


**Supplementary Figure. 1 LSSM shows great flexibility regarding wavelength choice.** (A) A brief illustration of the beam path of Zeiss Lightsheet.Z1. Samples are illuminated with a light-sheet. The emission light (fluorescence) and the scattered light is collected by the objective and then filtered by laser block filter. Emission selection modules consist of beam splitters and filters in front of the two cameras to collect different wavelengths of emission light simultaneously. (B) A collagen sample (2mg/ml) was tested in all 216 combination of lasers, laser block filters and emission selection modules. Laser power was set to 0.2%. Exposure time was 149.8 ms. Illumination mode: Single side. Pivot scan: On. Scale bars are 20 µm. Representative images from at least three independent experiments are shown.

**Movie legend**

**Movie1-3. Comparison of reconstructed 3D matrix structures obtained by LSSM or fluorescence.** Rat tail collagen (2 mg/ml) was fluorescently labeled with Atto 488. Z-stacks with step-size of 0.4 µm for 95 slices were obtained with LSSM and fluorescence mode concurrently at 37ºC. Reconstructed 3D matrix structures obtained by fluorescence modality is shown in Movie 1, by LSSM in Movie 2. Merged structured from both modalities is shown in Movie 3.

**Movie 4. Ultrathin filamentous structures observed with LSSM in SK-MEL-5 cells.** Rat tail collagen (2 mg/ml) was fluorescently labeled with Atto 488. Calcein red-orange loaded SK-MEL-5 cells were embedded in the matrix. Z-stacks with step-size of 0.82 µm for 52 slices were obtained using LSSM or fluorescence at 37ºC every 30 sec for 35.5 min. Time points during the occurrence of ultrathin filamentous structures is displayed slowly.

**Movie 5. Ultrathin filamentous structures observed with LSSM in 1.4E7 cells.** Rat tail collagen (2 mg/ml) was fluorescently labeled with Atto 488. Non-labeled 1.4E7 human pancreatic beta cells were embedded in the matrix. Z-stacks with step-size of 0.418 µm for 82 slices were obtained using LSSM or fluorescence at 37ºC every 20 sec for 10 min. The upper and lower frame highlights the ultrathin filamentous structures and the displacement of matrix networks by the cells, respectively.

**Movie 6. Visualization of a migrating T cell in ECM using LSSM.** Primary human $CD4^+$ T cells were loaded with CFSE and were embedded in the collagen matrices (Rat tail collagen, 2 mg/ml). Z-stacks with step-size of 1.5 μm for 70 slices were obtained using LSSM (gray) or fluorescence (green) at 37ºC every 1 min for 1 hour.

**Movie 7. Visualization of cell-induced displacement of collagen fibers.** Non-labeled SK-MEL-5 cells were embedded in rat tail collagen (2 mg/ml). Z-stacks with step-size of 1 μm for 295 slices were obtained using LSSM at 37ºC every 1 min for 2 hours. Pseudo colors are shown.
